# G-Protein-Coupled Receptor-Membrane Interactions Depend on the Receptor Activation state

**DOI:** 10.1101/743757

**Authors:** Apurba Bhattarai, Jinan Wang, Yinglong Miao

## Abstract

G-protein-coupled receptors (GPCRs) are the largest family of human membrane proteins and serve as primary targets of ∼1/3 of currently marketed drugs. In particular, adenosine A_1_ receptor (A_1_AR) is an important therapeutic target for treating cardiac ischemia-reperfusion injuries, neuropathic pain and renal diseases. As a prototypical GPCR, the A_1_AR is located within a phospholipid membrane bilayer and transmits cellular signals by changing between different conformational states. It is important to elucidate the lipid-protein interactions in order to understand the functional mechanism of GPCRs. Here, all-atom simulations using a robust Gaussian accelerated molecular dynamics (GaMD) method were performed on both the inactive (antagonist bound) and active (agonist and G protein bound) A_1_AR, which was embedded in a 1-palmitoyl-2-oleoyl-glycero-3-phosphocholine (POPC) lipid bilayer. In the GaMD simulations, the membrane lipids played a key role in stabilizing different conformational states of the A_1_AR. Our simulations further identified important regions of the receptor that interacted distinctly with the lipids in highly correlated manner. Activation of the A_1_AR led to differential dynamics in the upper and lower leaflets of the lipid bilayer. In summary, GaMD enhanced simulations have revealed strongly coupled dynamics of the GPCR and lipids that depend on the receptor activation state.

## Introduction

G-protein-coupled receptors (GPCRs) are primary cell surface receptors that account for vital physiological and pathological functions in human body. About 1/3 of currently marketed drugs approved by Food and Drug Administration (FDA) target GPCRs. Four subtypes of adenosine receptors (ARs), the A_1_AR, A_2A_AR, A_2B_AR and A_3_AR, mediate a broad range of physiological functions. Particularly, the Adenosine A_1_ Receptor (A_1_AR) has emerged as an important therapeutic target for treating cardiac ischemia-reperfusion injuries, neuropathic pain and renal diseases.(1) Being a GPCR, the A_1_AR is embedded in cell membrane, maintaining close contact with lipid molecules. Lipids have been suggested to affect the receptor conformation and dynamics, which play an important role in transmitting cellular signals from extracellular environment to the cytoplasm. Similarly, lipid metabolites are also known to bind proteins and act as messengers(2). These include lysolipids, sphingo-1-phosphate (S1P), diacylglycerol and fatty acyl derivates. In addition, lipids help in the partitioning of membrane and receptors. Membrane proteins are affected by lipid compositions and function differently in healthy and diseased individuals.(3) Therefore, it is important to study GPCR-membrane interactions in order to elucidate functional mechanism of the membrane proteins.

Experimental techniques including fluorescence resonance energy transfer (FRET), fluorescence correlation spectroscopy (FCS), fluorescence recovery after photobleaching (FRAP) and fluorescence-based monitoring of solvent relaxations rates have been utilized to study protein-membrane biology(4–7). Experiments showed that cholesterol could affect the stability, oligomerization, and ligand binding properties of membrane proteins(8–21). X-ray crystal structures identified allosteric sites for cholesterol binding to GPCRs(13, 15, 22). Phospholipids were found to modulate dynamic processes of GPCRs such as the G protein association and ligand binding(23, 24). Recently, Dawaliby et. al. showed experimentally that lipids with different head groups favor different activation states of the β_2_-adrenergic receptors (β_2_AR)(25). Lipids with phosphatidylglycerol (PG) headgroups preferred agonist binding and receptor activation, whereas lipids with phosphatidylethanolamine (PE) headgroups preferred antagonist binding and inactive state of the β_2_AR. Despite these advances, there remains a knowledge gap in the understanding of protein-membrane interactions. From atomic motions of lipid molecules to curvature change across the cell membrane, protein-membrane interactions span a wide range of time scales(26). It is often difficult to directly examine protein-membrane interactions in experiments due to limited time resolution.

Molecular dynamics (MD) simulation has emerged as a powerful computational technique to bridge the gap of knowledge for studying membrane-protein interactions. Both atomistic and coarse-grained MD simulations have been applied to study the effects of lipids in protein dynamics and cellular signaling.(3) Bruzzese et al. confirmed the above mentioned experimental results obtained by Dawaliby et. al. that different charges of PG and PE lipid headgroups affected the GPCR activation and deactivation in MD simulations(27). The negative charge in PG molecules favored interaction with positively-charged residues in the intracellular loop 3 (ICL3) and intracellular end of transmembrane helix 6 (TM6). This stabilized the outward movement of TM6 and hence the active state of the β_2_AR. Neale et. al.(28) showed that the PG lipid blocked formation of the R131^3.50^-E268^6.30^ ionic lock by interacting with R131^3.50^ in the β_2_AR. Residue superscripts denote Ballesteros and Weinstein (BW) numbering of GPCRs(29). The R^3.50^ and E^6.30^ are highly conserved residues in GPCRs and often form an ionic lock in inactive receptors. In contrast, lipids with negatively-charged PE headgroup formed favorable interactions with the positively-charged residues in the TM6 and stabilized active state of the β_2_AR(27). Salas-Estrada et. al(30) showed that activation in rhodopsin induced changes in the membrane structure, including increase in the local order and effective length of lipid acyl chains in the vicinity of the protein. Dror et. al. showed that gradual inactivation of the β_2_AR occurred in the neutral lipid 1-palmitoyl-2-oleoyl-glycero-3-phosphocholine (POPC)(31, 32). This was consistent with another study favored partial deactivation of the β_2_AR was found in the 1,2-dioleoyl-*sn*-glycero-3-phosphocholine (DOPC) lipids(27). In coarse-grained MD simulations, Song et. al. showed that PIP_2_ (Phosphatidylinositol 4,5-bisphosphate) stabilized the outward movement of TM6 by binding in the crevice between TM6 and TM7 of adenosine A_2A_ receptor (A_2A_AR)(33).

In addition, enhanced sampling techniques have been applied to investigate protein membrane interaction. Using steered MD(34) and umbrella sampling(35) techniques, Song et. al. suggested that the PIP_2_ facilitated recruitment of the G protein by forming bridging interactions with basic residues of the G_*α*_ subunit and hence stabilizing the active A_2A_AR(33). However, these enhanced sampling techniques require predefined collective variables, which are often difficult to identify in the context of protein-membrane interactions. In this context, Gaussian accelerated MD (GaMD) is a robust technique that provides unconstrained enhanced sampling without the need to set predefined collective variables(36, 37). GaMD simulations have been successfully applied to investigate GPCR activation(36, 38, 39), protein folding(36, 39), ligand binding and unbinding(36, 38, 39), protein-protein interactions(40–42) and protein-nucleic acid interactions(43, 44).

Here, we have applied GaMD to investigate lipid interactions with the A_1_AR in two different conformational states, the cryo-EM structure of the active adenosine ADO-bound A_1_AR coupled with the G_i_ protein (referred to as ADO-A_1_AR-Gi)(45) and the X-ray structure(46, 47) of the inactive antagonist PSB36-bound A_1_AR (referred to as PSB36-A_1_AR). Our simulations showed that the protein-membrane interactions depended on different conformational states of the A_1_AR. The membrane lipids played an important role in stabilizing different conformations of the A_1_AR. GaMD simulations further identified important regions of the receptor that interacted distinctly with the lipids. Activation of the A_1_AR led to differential dynamics in the upper and lower leaflets of the lipid bilayer.

## Materials and Methods

### Gaussian Accelerated Molecular Dynamics (GaMD)

GaMD is an enhanced sampling technique, in which a harmonic boost potential is added to reduce the system energy barriers(36). GaMD is able to accelerate biomolecular simulations by orders of magnitude(39, 48). GaMD does not need predefined collective variables. Moreover, because GaMD boost potential follows a gaussian distribution, biomolecular free energy profiles can be properly recovered through cumulant expansion to the second order(36). GaMD has successfully overcome the energetic reweighting problem in free energy calculations that was encountered in the previous amd method(49, 50) for free energy calculations. GaMD has been implemented in widely used software packages including AMBER (36, 51) and NAMD(52). A brief summary of GaMD is provided here.

Consider a system with *N* atoms at positions 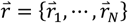. When the system potential 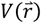 is lower than a reference energy *E*, the modified potential 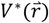 of the system is calculated as:

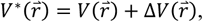

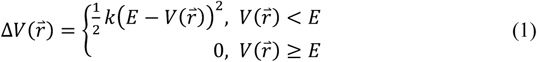

where *k* is the harmonic force constant. The two adjustable parameters *E* and *k* are automatically determined based on three enhanced sampling principles(36). The reference energy needs to be set in the following range:

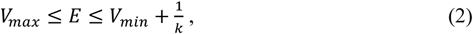

where *V*_*max*_ and *V*_*min*_ are the system minimum and maximum potential energies. To ensure that Eqn. (2) is valid, *k* has to satisfy: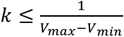 Let us define 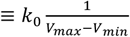, then 0 < *k*_0_ ≤ 1. The standard deviation of Δ*V* needs to be small enough (i.e., narrow distribution) to ensure proper energetic reweighting(53): *σ*_Δ*V*_ = *k*(*E* – *V*_*avg*_) *σ*_*V*_ ≤ *σ*_0_ where *V*_*avg*_ and *σ*_*V*_ are the average and standard deviation of the system potential energies, *σ*_Δ*V*_ is the standard deviation of Δ*V* with *σ*_0_ as a user-specified upper limit (e.g., 10*k*_*B*_T) for proper reweighting. When *E* is set to the lower bound *E=V*_*max*_, *k*_0_ can be calculated as:

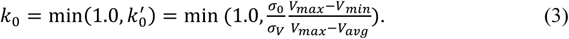

Alternatively, when the threshold energy *E* is set to its upper bound 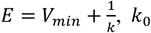 is set to:

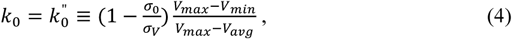

if 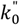 is found to be between *0* and *1*. Otherwise, *k*_0_ is calculated using Eqn. (3).

Similar to aMD, GaMD provides schemes to add only the total potential boost Δ*V*_*P*_, only dihedral potential boost Δ*V*_*D*_, or the dual potential boost (both Δ*V*_*P*_ and Δ*V*_*D*_). The dual-boost simulation generally provides higher acceleration than the other two types of simulations(54). The simulation parameters comprise of the threshold energy E for applying boost potential and the effective harmonic force constants, *k*_0*P*_ and *k*_0*D*_ for the total and dihedral potential boost, respectively.

### Energetic Reweighting of GaMD Simulations

To calculate potential of mean force (PMF)(55) from GaMD simulations, the probability distribution along a reaction coordinate is written as *p*\*(*A*). Given the boost potential 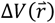 of each frame, *p*\*(*A*) can be reweighted to recover the canonical ensemble distribution, *p*(*A*), as:

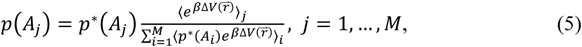

where *M* is the number of bins, β = *k*_*B*_*T* and 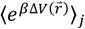 is the ensemble-averaged Boltzmann factor of 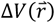 for simulation frames found in the *j*^th^ bin. The ensemble-averaged reweighting factor can be approximated using cumulant expansion:

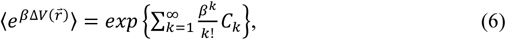

where the first two cumulants are given by

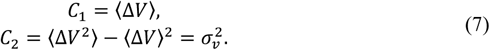

The boost potential obtained from GaMD simulations usually follows near-Gaussian distribution. Cumulant expansion to the second order thus provides a good approximation for computing the reweighting factor(36, 53). The reweighted free energy *F*(*A*) = -*k*_*B*_*T* ln *p*(*A*) is calculated as:

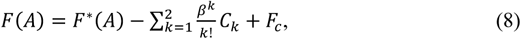

where *F*\*(*A*) = -*k*_*B*_*T* ln *p*\*(*A*) is the modified free energy obtained from GaMD simulation and *F*_*c*_ is a constant.

### Lipid -S_CD_ Order Parameter

The -S_CD_ order parameter measures orientational anisotropy of the C-H bond in sn-2 acyl chains of lipids that are usually obtained from NMR experiments. It is a function of the angle between the C-H bond and lipid bilayer normal. It is defined by following equation:

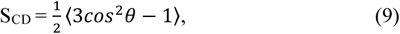

where *θ* is the angle between the bilayer normal and C-H bond and ⟨…⟩ denotes an ensemble average. Here, the -S_CD_ order parameter is averaged over all the lipids in the system and all the frames in the simulation trajectory. The -S_CD_ order parameter calculated from GaMD simulations is not reweighted due to complexity of the function. However, since GaMD maintained the overall shape of the original potential energy surface(36), the resulting order parameter is found to be close to the experimental values(56) (see **Results**). The -S_CD_ value usually ranges from -0.25 to 0.5, with 0.5 for the C-H bond being fully ordered along the bilayer normal and -0.25 being parallel to the bilayer plane. The -S_CD_ approximates the mobility of each C-H bond and hence estimates the membrane fluidity.

### System Setup

The cryo-EM structure of the ADO-A_1_AR-Gi complex (PDB: 6D9H(45)) and X-ray structure of PSB36-A_1_AR complex (PDB: 5N2S(46)) were used to prepare the simulation systems. As helix 8 region was missing in the crystal structure of PSB36-A_1_AR, atomic coordinates were added using another X-ray structure of the inactive A_1_AR (PDB: 5UEN(47)) after aligning the receptor TM domain. All chain termini were capped with neutral groups, i.e. the acetyl group (ACE) for the N-terminus and methyl amide group (CT3) for C terminus. Protein residues were set to the standard CHARMM protonation states at neutral pH with the *psfgen* plugin in VMD(57). Then the receptor was inserted into a POPC bilayer with all overlapping lipid molecules removed using the *Membrane* plugin in VMD(57). The system charges were then neutralized at 0.15 M NaCl using the *Solvate* plugin in VMD(57). Periodic boundary conditions were applied on the simulation systems. The simulation systems of the active and inactive A_1_AR systems are summarized in **Table 1**.

**Table 1:**
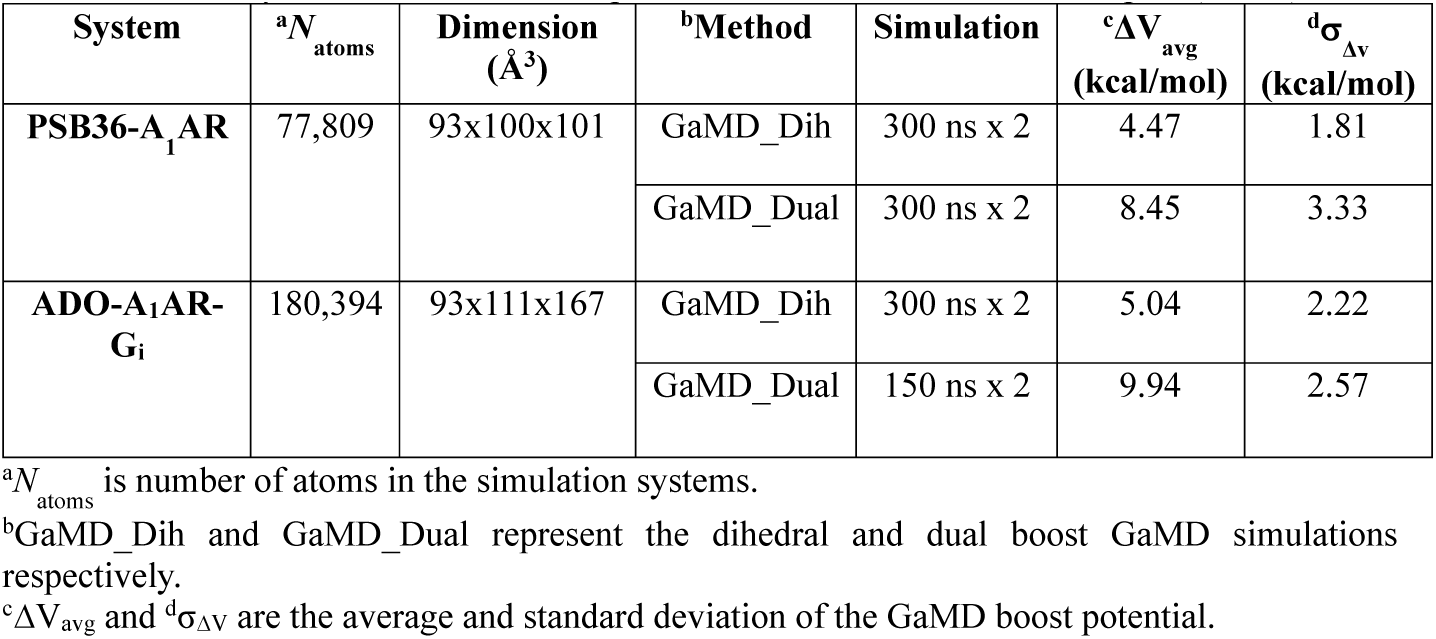
Summary of GaMD simulations performed on the adenosine A_1_ receptor (A_1_AR).

| **System** | **<sup>a</sup>N<sub>atoms</sub>** | **Dimension**<br>**(Å<sup>3</sup>)** | **<sup>b</sup>Method** | **Simulation** | **<sup>c</sup>ΔV<sub>avg</sub>**<br>**(kcal/mol)** | **<sup>d</sup>σ<sub>Δv</sub>**<br>**(kcal/mol)** |
| --- | --- | --- | --- | --- | --- | --- |
| **PSB36-A<sub>1</sub>AR** | 77,809 | 93x100x101 | GaMD_Dih | 300 ns x 2 | 4.47 | 1.81 |
|  |  |  | GaMD_Dual | 300 ns x 2 | 8.45 | 3.33 |
| **ADO-A<sub>1</sub>AR-G<sub>i</sub>** | 180,394 | 93x111x167 | GaMD_Dih | 300 ns x 2 | 5.04 | 2.22 |
|  |  |  | GaMD_Dual | 150 ns x 2 | 9.94 | 2.57 |

<sup>a</sup>N<sub>atoms</sub> is number of atoms in the simulation systems.
<sup>b</sup>GaMD\_Dih and GaMD\_Dual represent the dihedral and dual boost GaMD simulations respectively.
<sup>c</sup>ΔV<sub>avg</sub> and <sup>d</sup>σ<sub>Δv</sub> are the average and standard deviation of the GaMD boost potential.

### Simulation Protocol

The CHARMM36 parameter set(58) was used for the protein and POPC lipids. For agonist ADO and antagonist PSB36, the force field parameters were obtained from the CHARMM ParamChem web server(59, 60). Initial energy minimization and thermalization of the A_1_AR system follow the same protocol as used in the previous GPCR simulations(61). The simulation proceeded with equilibration of lipid tails. With all the other atom fixed, the lipid tails were energy minimized for 1000 steps using the conjugate gradient algorithm and melted with constant number, volume, and temperature (NVT) run for 0.5ns at 310 K. Each system was further equilibrated using constant number, pressure, and temperature (NPT) run at 1 atm and 310 K for 10 ns with 5 kcal (mol Å^2^)^-1^ harmonic position restraints applied to the protein. Further equilibration of the systems was performed using an NPT run at 1 atm and 310 K for 0.5ns with all atoms unrestrained. Conventional MD simulation was performed on each system for 10 ns at 1atm pressure and 310 K with a constant ratio constraint applied on the lipid bilayer in the X-Y plane. The GaMD simulations were carried out using NAMD2.13(52, 62). Both dihedral and dual-boost GaMD simulations were then performed to study the protein-membrane interactions in the inactive and active A_1_AR systems (**Table 1**). In the GaMD simulations, the threshold energy *E* for adding boost potential is set to the lower bound, i.e. *E = V*_max_(36, 52). The simulations included 50ns equilibration after adding the boost potential and then multiple independent production runs lasting 150 – 300 *ns* with randomized initial atomic velocities. GaMD production simulation frames were saved every 0.2ps for analysis.

### Simulation analysis

The VMD(57) and CPPTRAJ(63) tools were used for trajectory analysis. In particular, distance was calculated between the Cα atoms residues Arg^3.50^ and Glu^6.30^. Root-mean-square fluctuations (RMSFs) were calculated for the protein residues and ligands, averaged over two independent GaMD simulations and color coded for schematic representation of each complex system. MEMBPLUGIN, a plugin for the VMD package was used to calculate the -S_CD_ order parameter for POPC lipid tails(64). The -S_CD_ order parameters were averaged over all lipids and frames of the two independent GaMD simulations for each system. The CPPTRAJ tool was used to calculate the correlation matrices. The Cα atoms of the receptor and phosphorous atoms in the POPC lipid head groups were used for the calculations. In addition to the phosphorous atom, the C_8_ and C_18_ atoms representing different regions of the lipids were also used to calculate dynamic correlations with the receptor. The *PyReweighting* toolkit(53) was applied to reweight GaMD simulations for free energy calculations by combining independent trajectories for each system. A bin size of 1 Å was used for the Arg^3.50^-Glu^6.30^ distance and 1 for the number of lipids. The cutoff was set to 500 for calculating the 2D PMF profiles.

## Results

### Structural flexibility of the A_1_AR depended on the receptor conformational state

All-atom GaMD simulations were performed on two different conformational states of the A_1_AR, active (ADO-A_1_AR-Gi) and inactive (PSB36-A_1_AR) states (**Table 1**). For the inactive A_1_AR system, the boost potential was 4.47±1.81 kcal/mol and 8.45±3.33 kcal/mol in dihedral and dual-boost GaMD simulations, respectively. For the active A_1_AR system, the boost potential was 5.04±2.22 kcal/mol and 9.94±2.57 kcal/mol in dihedral and dual-boost GaMD simulations, respectively (**Table 1**). Thus, dual-boost GaMD provided higher acceleration in the simulations with greater boost potential. In the dihedral GaMD simulations of the inactive A_1_AR, the TM helices of the receptor were rather rigid. Only the intracellular end of TM6, the terminus of helix 8 (H8), extracellular end of TM1 and extracellular loop 2 (ECL2) regions were flexible (**Figure 1A**). Similar results were obtained for the active A_1_AR in the dihedral GaMD simulations (**Figure 1B**). However, the intracellular ends of TM6 and TM5 of the A_1_AR exhibited more fluctuations in the active state compared to the inactive state. The ECL2 region was relatively more flexible in the active A_1_AR than in the inactive A_1_AR. In both systems, the ligands remained stably bound at the orthosteric site throughout the simulations. In comparison, the G protein coupled to the active A_1_AR exhibited higher fluctuations. In particular, C terminus of the α5 helix, α4-β5 loop and α4-β6 loop of the Gα subunit and terminal ends of the G_βγ_ subunits exhibited high fluctuations up to 3 Å. Similar results were also found in the dual-boost GaMD simulations of the inactive and active A_1_AR systems (**Figure S1**).

**Figure 1:**
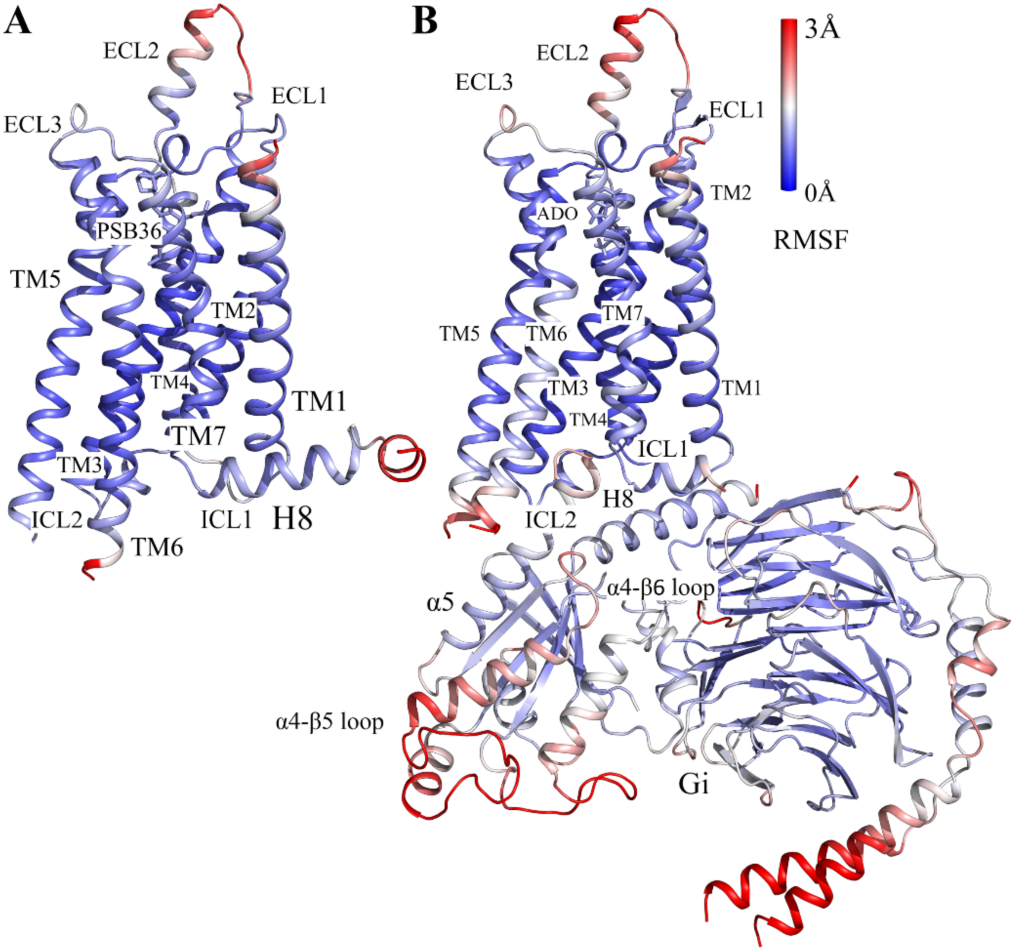
Comparison of structural flexibility of the inactive and active A1AR systems obtained from dihedral GaMD simulations: (A) Root-mean-square fluctuations (RMSFs) of the inactive PSB36-A1AR complex. (B) RMSFs of the active ADO-A1AR-Gi protein complex. A color scale of 0 Å (blue) to 3 Å (red) was used.

### Lipids in the lower leaflet of the active A_1_AR system showed higher fluidity than in the inactive-A_1_AR system

The lipid -S_CD_ order parameters were calculated for the upper (extracellular) and lower (cytoplasmic) leaflets from GaMD simulations of the inactive and active A_1_AR systems (**Figure 2**). In both inactive and active A_1_AR systems, lower leaflet was more fluid than the upper leaflet with smaller -S_CD_ order parameters. In the upper leaflet, the -S_CD_ order parameter of POPC lipid molecules were comparable between the inactive and active systems. In contrast, the lower leaflet exhibited significant differences between the inactive and active A_1_AR systems. In particular, the -S_CD_ order parameter of the fifth carbon atom in POPC was ∼0.20 in the lower leaflet in the inactive A_1_AR system (**Figure 2A, 2C**), but decreased to ∼0.17 in the active A_1_AR system (**Figure 2B, 2D**). This indicated higher inclination of C-H bonds being ordered along the bilayer normal in the lower leaflet of the active A_1_AR system. This appeared to correlate with the outward movement of TM6 as the A_1_AR changed from the inactive to active state. In the inactive A_1_AR, the R131^3.50^-E268^6.30^ distance at the free energy minimum was ∼7 Å (**Figure 3A, 3C**). In comparison, this distance at the free energy minimum increased to ∼17 Å in the active A_1_AR (**Figure 3B, 3D**). The lateral movement of the TM6 could push the surrounding lipids. Higher flexibility of the receptor TM6 intracellular end was accompanied by increased fluidity of lipids in the lower leaflet of membrane. Similar results were found from the dihedral and dual-boost GaMD simulations (**Figure 3**). Therefore, the structural conformation and flexibility of the GPCR are strongly coupled with the surrounding membrane lipids.

**Figure 2:**
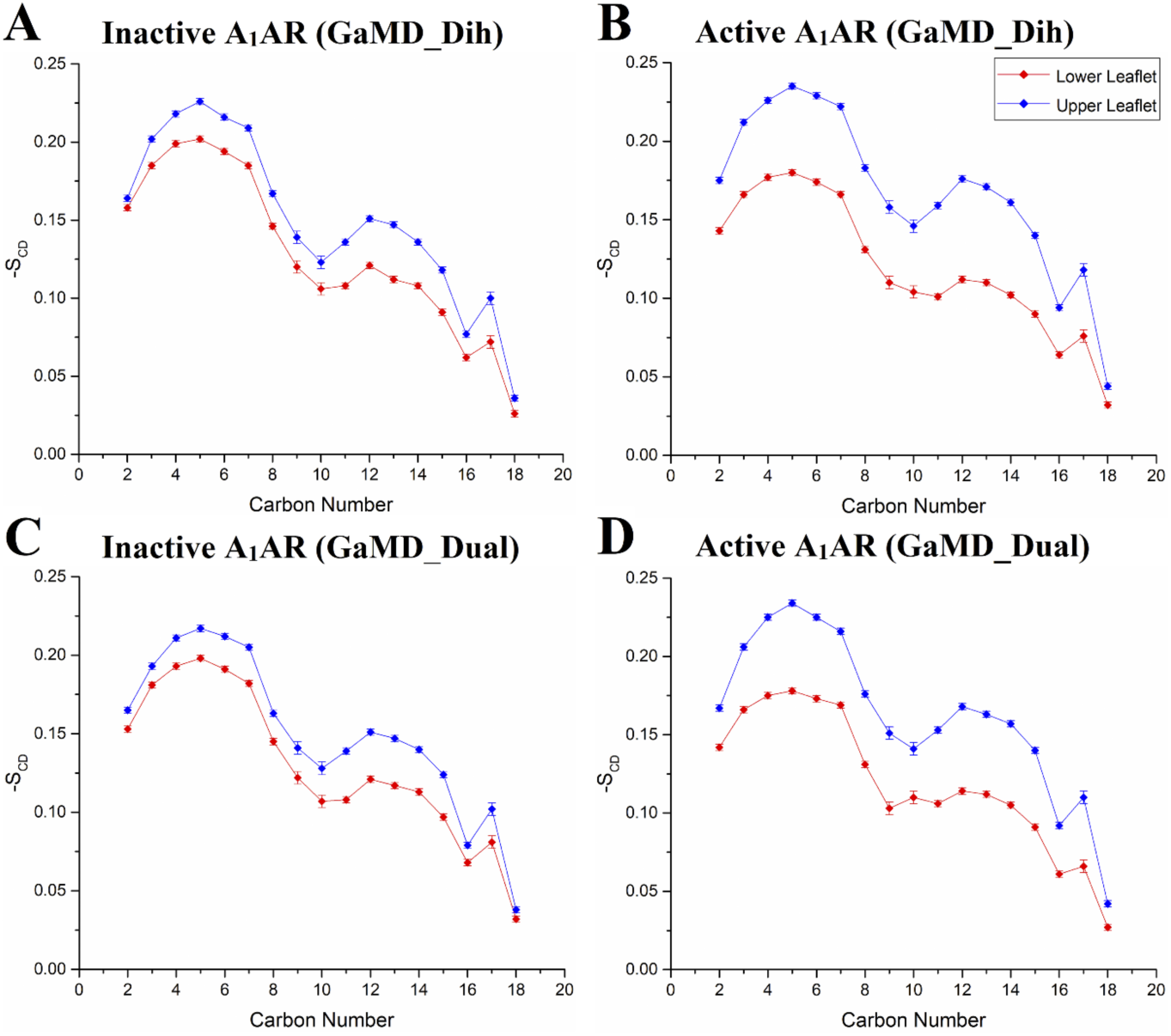
The -S_CD_ order parameters calculated for sn-2 acyl chains of POPC lipids in different simulation systems: (A) Inactive A1AR using dihedral-boost GaMD, (B) Active A1AR using dihedral-boost GaMD, (C) Inactive A1AR using dual-boost GaMD and (D) Active A1AR using dual-boost GaMD. Red diamond lines represent the average -S_CD_ order parameters for the cytoplasmic lower leaflet and blue diamond lines for the extracellular upper leaflet.

**Figure 3:**
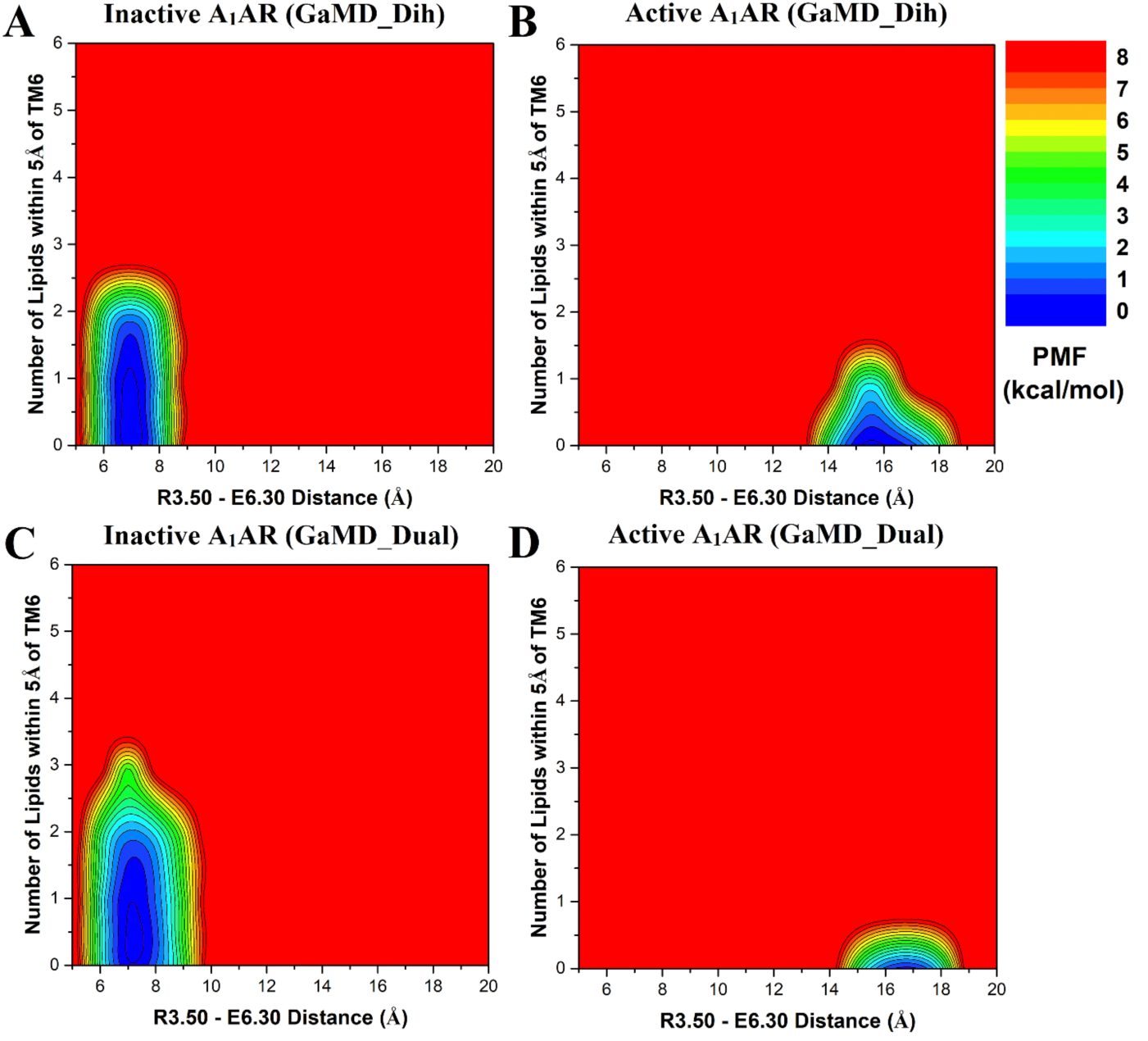
Free energy profiles of the extracellular upper leaflet of membrane in different simulation systems regarding the number of lipids within 5 Å of the receptor TM6 and the receptor R3.50 – E6.30 distance: (A) Inactive A_1_AR using dihedral-boost GaMD, (B) Active A_1_AR using dihedral-boost GaMD, (C) Inactive A_1_AR using dual-boost GaMD and (D) Active A_1_AR using dual-boost GaMD. The R3.50 – E6.30 distance is ∼7 Å in the inactive A1AR and increases to ∼17 Å in the active A1AR due to outward movement of TM6.

### The inactive A_1_AR attracted more lipids in the upper leaflet to the TM6 than the active A_1_AR

Considering significant conformational changes (especially in the TM6) during receptor activation, we hypothesized that the number of lipids interacting with the active and inactive receptor are different. In order to identify the low-energy states of the membrane-receptor interactions, potential of mean force (PMF) profiles were calculated by reweighting the GaMD simulations (**Figure 3**). The R131^3.50^-E268^6.30^ distance was chosen as one reaction coordinate to characterize activation of the GPCR. The number of POPC phosphate head groups within 5 Å of TM6 was calculated as the other reaction coordinate. In the upper leaflet, approximately one lipid molecule was found interacting with TM6 in the inactive A_1_AR (**Figure 3A, 3C**). But no lipid in the upper leaflet was found within to 5 Å of TM6 in the active A_1_AR (**Figure 3B, 3D**). Further analysis revealed that one positively-charged residue was located in the receptor ECL3 (K265^ECL3^), being close to the extracellular end of the TM6 (**Figure 5A**). In the inactive A_1_AR, this lysine pointed towards the lipid membrane and thus attracted the negatively-charged phosphate head group of a POPC molecule. Instead, the positively-charged side chain of K265^ECL3^ formed a stable salt-bridge with negatively-charged glutamate (E172^ECL2^) of ECL2 in the active A_1_AR (**Figure S4**). Residue K265^ECL3^ did not interact with the lipid in the active A_1_AR. Therefore, the inactive A_1_AR interacted with more phospholipids in the upper leaflet compared to the active A_1_AR.

### The active A_1_AR attracted more lipids in the lower leaflet to the TM6 than the inactive A_1_AR

In contrast to the upper leaflet, the lower leaflet had more lipids within 5 Å of TM6 in the active A_1_AR than in the inactive system (**Figure 4**). In the lowest energy state, the inactive A_1_AR interacted with approximately two lipids within 5 Å of TM6 (**Figure 4A, 4C**). In comparison, the active A_1_AR exhibited a relatively broader energy well. The TM6 intracellular domain interacted with ∼2-4 lipid molecules (**Figure 4B, 4D**). Upon activation of the A_1_AR, the TM6 moved outwards by ∼10 Å and exposed its positively-charged residues to the membrane. Therefore, the lipids that diffused in the lower leaflet of the membrane interacted more frequently with the receptor. Fewer lipids interacted with the inactive A_1_AR because the receptor has a narrower curvature in the TM6 intracellular region. Moreover, the lower leaflet had a larger number of lipids interacting with receptor TM6 than the upper leaflet. Out of five positively-charged residues in TM6, four were located in the intracellular region (K224^6.25^, K228^6.29^, K231^6.32^ and K234^6.35^) (**Figure 5B**). The negatively-charged phosphate head groups of POPC tended to interact with these positively-charged lysine residues to stabilize the active receptor conformation. Therefore, the lipid-GPCR interaction should play an important role in the conformational changes during activation of the A_1_AR.

**Figure 4:**
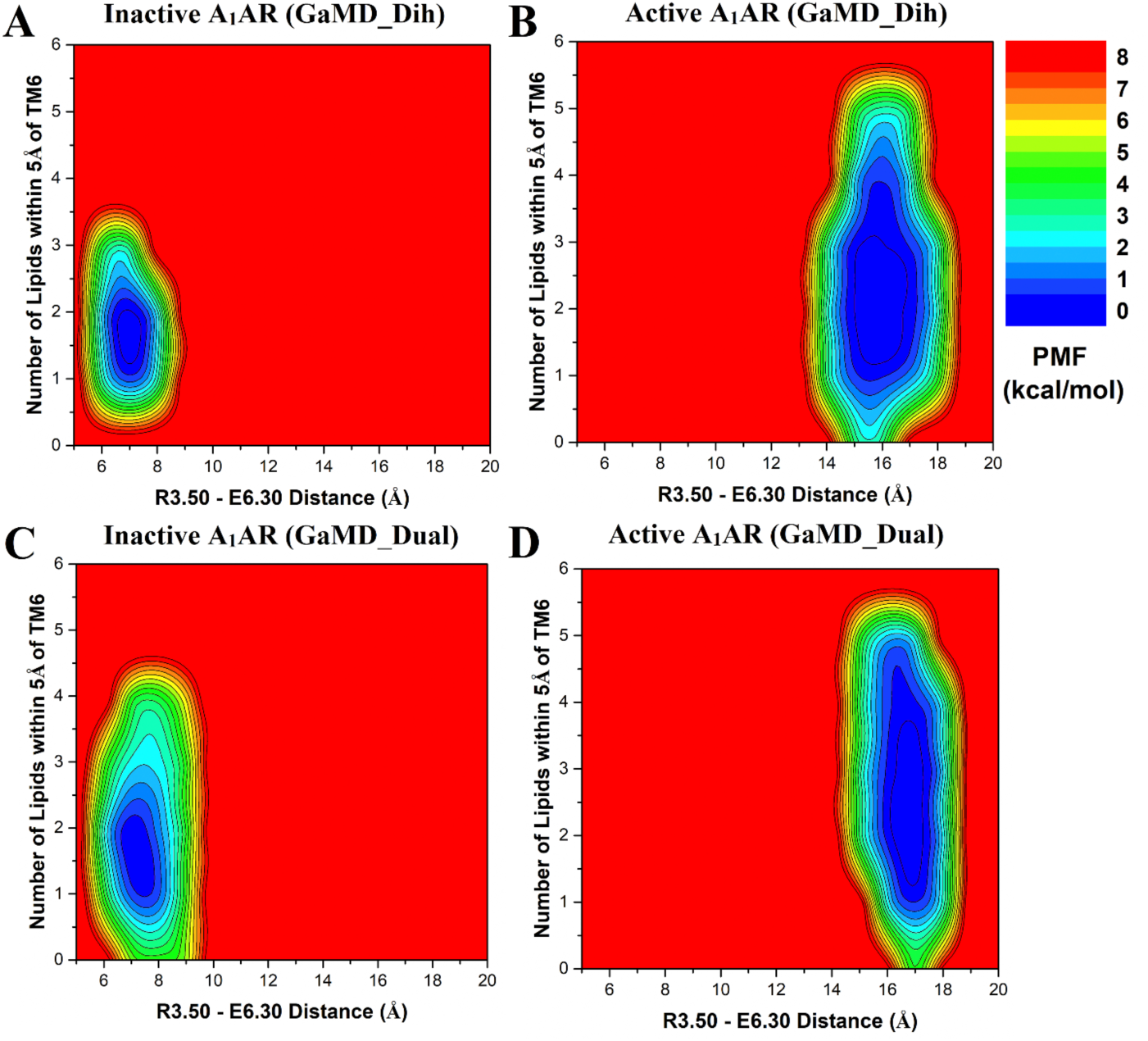
Free energy profiles of the cytoplasmic lower leaflet of membrane in different simulation systems regarding the number of lipids within 5 Å of the receptor TM6 and the receptor R3.50 – E6.30 distance: (A) Inactive A_1_AR using dihedral-boost GaMD, (B) Active A_1_AR using dihedral-boost GaMD, (C) Inactive A_1_AR using dual-boost GaMD and (D) Active A_1_AR using dual-boost GaMD. The R3.50 – E6.30 distance is ∼7 Å in the inactive A1AR and increases to ∼17 Å in the active A1AR due to outward movement of TM6.

**Figure 5:**
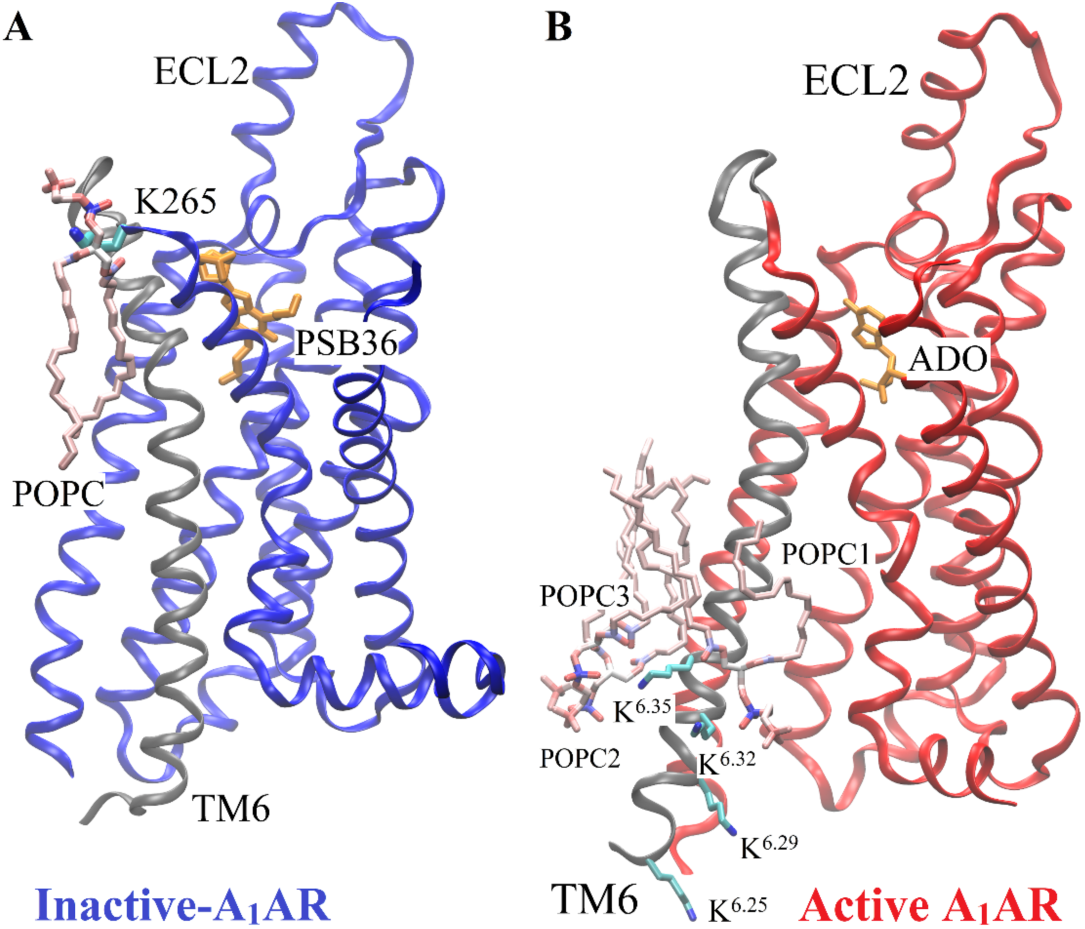
Minimum energy states of POPC lipid interacting with the positively-charged lysine residues in TM6 of the receptor obtained from dihedral GaMD simulations. (A) One POPC molecule in the upper leaflet interacts with one lysine residue (K265^ECL3^) of the inactive A_1_AR. (B) Three POPC molecules (POPC1, POPC2, POPC3) in the lower leaflet interact with four Lysine residues (K263^6.25^, K267^6.29^, K270^6.32^ and K273^6.35^) of the active A_1_AR. The receptor TM6 is colored in gray.

### GaMD simulations revealed strongly coupled dynamics between the GPCR and membrane lipids

Dynamic correlations were identified between residues of the A_1_AR and lipids in both the upper and lower leaflets. The C*α* atoms in the receptor residues and the phosphorous atoms in the lipid head groups were used to calculate the correlation matrices (see details in **Methods**). Similar results were obtained using the C_8_ and C_18_ atoms in the lipid hydrophobic tails to calculate the dynamic correlation matrices (**Figure S5**). In all the simulation systems, motions of the receptor N-terminus, ECL1, ECL2 and ECL3 regions were positively correlated to those of lipids in upper leaflet (**Figure 6**). Similarly, motions of the receptor ICL1, ICL2 and ICL3 were positively correlated to those of lipids in the lower leaflet (**Figure 7**). For the inactive A_1_AR, most TM helix residues exhibited negatively correlated motions with the lipids. In this regard, the TM helices appeared to move in the opposite direction relative to the lipids in the inactive A_1_AR simulation system (**Figure 6A, 6C, 7A and 7A**). For the active A_1_AR, correlations between TM helix residues and the lipids in both the upper and lower leaflets were very weak, being close to zero (**Figure 6B, 6D, 7B and 7D**). However, marked positive correlations were identified between the intracellular region of the receptor TM6 and lipids in the lower leaflet (**Figures 7B and 7D**). Therefore, the TM6 intracellular region of the active A_1_AR appeared to move in the same direction with the surrounding lipids. This was highly consistent with the simulation finding that significantly more lipids were found within 5 Å of the TM6 intracellular domain in the active A_1_AR system (**Figure 4**) and they formed remarkably stronger electrostatic interactions (**Figure 5**) compared with the inactive A_1_AR system. In summary, the GaMD simulations revealed strongly coupled dynamics between the GPCR and membrane lipids.

**Figure 6:**
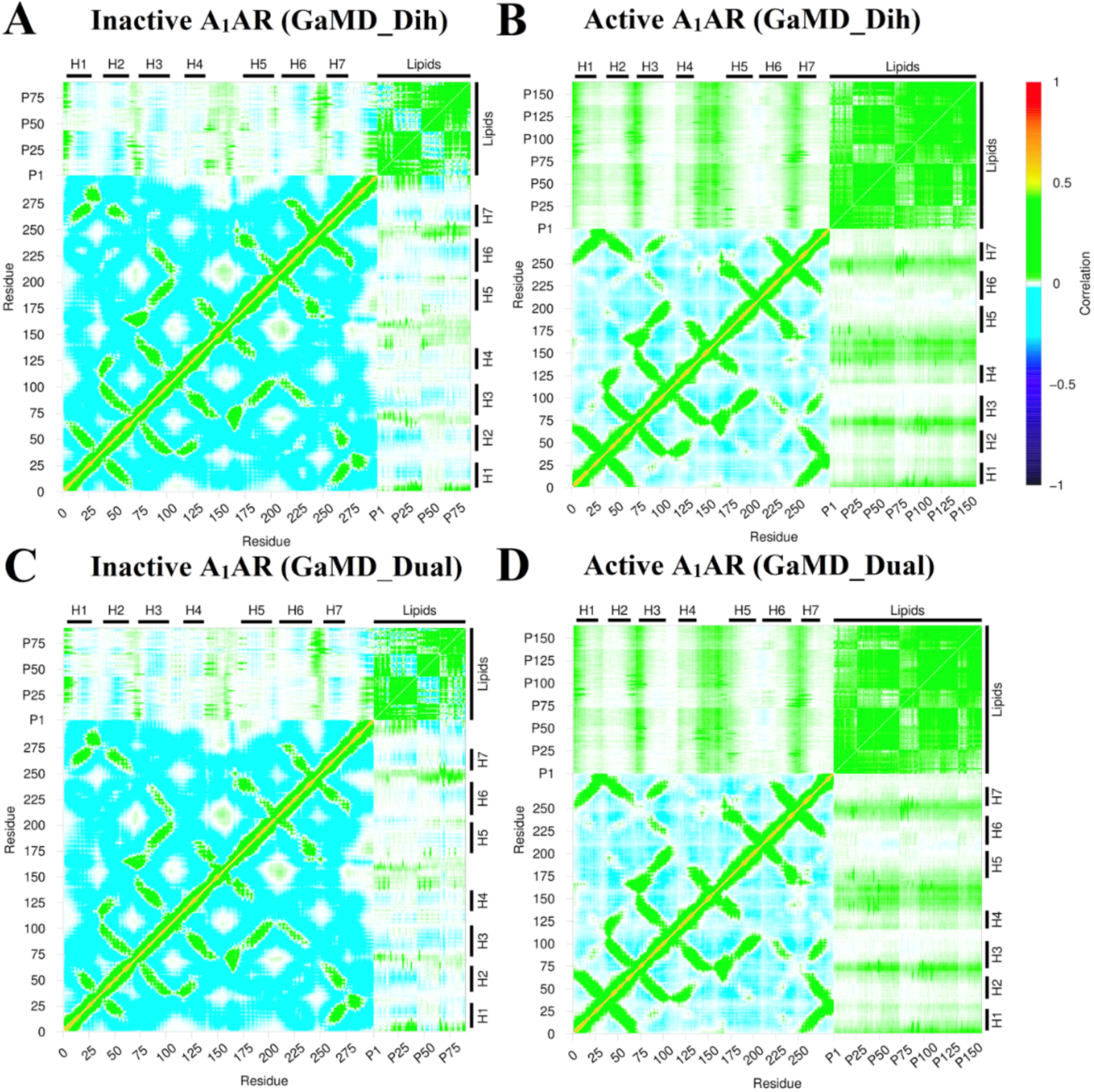
Dynamic correlation matrices calculated for lipids in the extracellular upper leaflet with residues in the A1AR in different simulation systems: (A) Inactive A_1_AR using dihedral-boost GaMD, (B) Active A_1_AR using dihedral-boost GaMD, (C) Inactive A_1_AR using dual-boost GaMD and (D) Active A_1_AR using dual-boost GaMD. The C*α* atoms of the receptor and phosphorous atoms in the lipid head groups were used for calculating the correlation matrices here. Similar results were obtained using the C_8_ and C_18_ atoms in the lipid hydrophobic tails as shown in **Figure S5**. The receptor ICL1, ICL2 and ICL3 represent intracellular loops between TM helices 1-2, 3-4, and 5-6 respectively. Similarly, the receptor ECL1, ECL2 and ECL3 represent extracellular loops between TM helices 2-3, 4-5, and 6-7 respectively.

**Figure 7:**
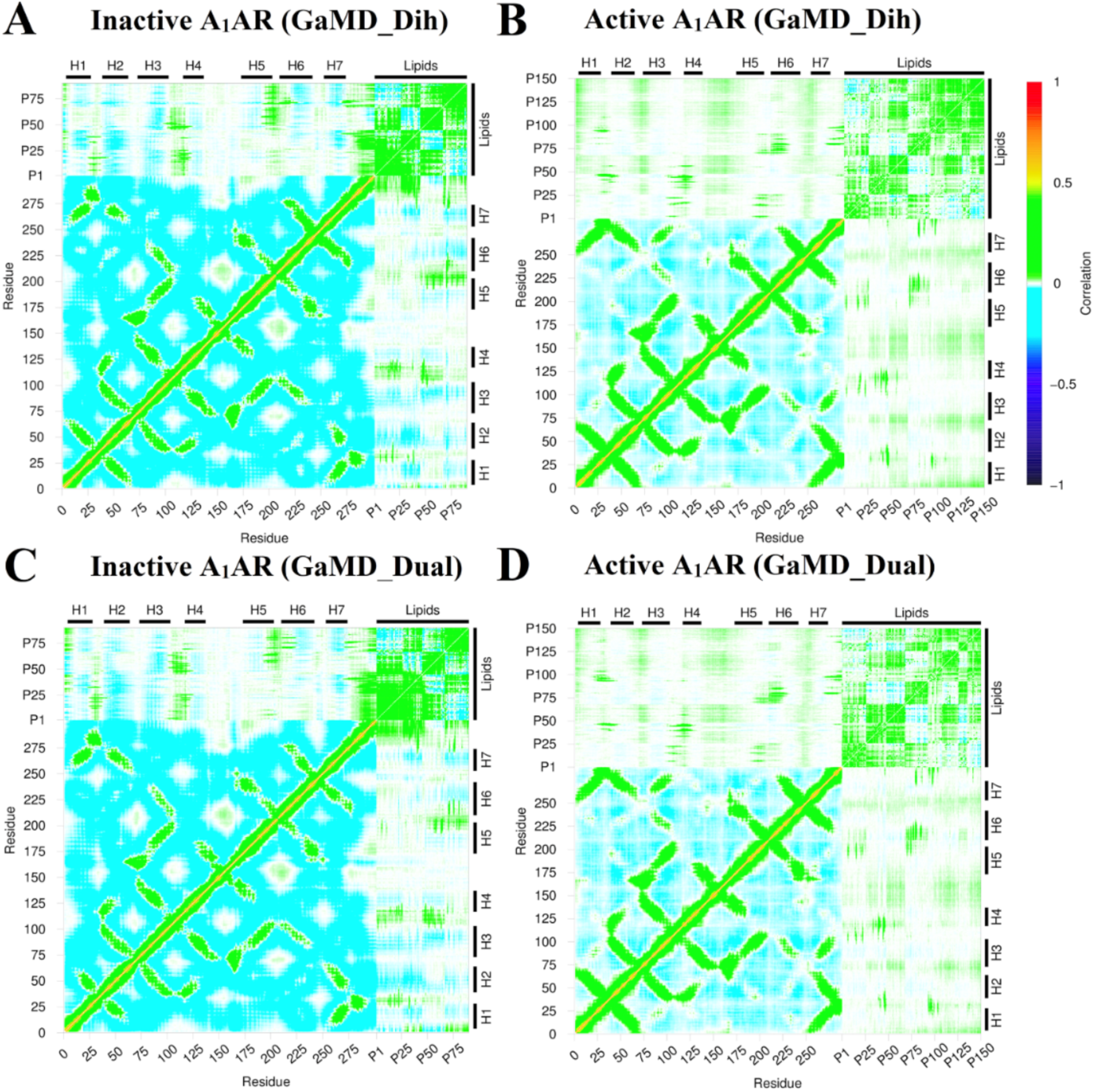
Dynamic correlation matrices calculated for lipids in the intracellular lower leaflet with residues in the A1AR in different simulation systems: (A) Inactive A_1_AR using dihedral-boost GaMD, (B) Active A_1_AR using dihedral-boost GaMD, (C) Inactive A_1_AR using dual-boost GaMD and (D) Active A_1_AR using dual-boost GaMD. The C*α* atoms of the receptor and phosphorous atoms in the lipid head groups were used for calculating the correlation matrices here. Similar results were obtained using the C_8_ and C_18_ atoms in the lipid hydrophobic tails as shown in **Figure S5**. The receptor ICL1, ICL2 and ICL3 represent intracellular loops between TM helices 1-2, 3-4, and 5-6 respectively. Similarly, the receptor ECL1, ECL2 and ECL3 represent extracellular loops between TM helices 2-3, 4-5, and 6-7 respectively.

## Discussion

In this study, we have applied all-atom GaMD simulations to investigate GPCR-membrane interactions, using the A_1_AR as a model receptor. In the GaMD simulations, the inactive and active A_1_AR showed different structural flexibility profiles. The ECL2 region, intracellular ends of TM6 and TM5 exhibited higher fluctuations in the active A_1_AR compared to the inactive A_1_AR. The receptor TM domain was rigid and the ligands remained tightly bound at the orthosteric site. However, the G protein coupled to the active A_1_AR exhibited high flexibility during the simulations, especially in the α5 helix, α4-β5 loop and α4-β6 loop of the Gα subunit and terminal ends of the G_βγ_ subunits. These results were consistent to our earlier simulation findings of the AR-G protein complexes(65).

The -S_CD_ order parameter values obtained from GaMD simulations were consistent with experimental data. In NMR experiments, the -S_CD_ order parameter for the fifth carbon C-H bond of POPC was observed to be at ∼0.18-0.20(56). The -S_CD_ order parameter of POPC’s fifth carbon atom was ∼0.20±0.02 in the lower leaflet in the inactive A_1_AR system. It decreased to ∼0.17±0.02 in the inactive A_1_AR system. The -S_CD_ order parameter of the ninth carbon C-H bond in POPC was measured as ∼0.10 in NMR experiments(56), for which the same value was obtained from GaMD simulations. Furthermore, the GaMD simulations showed that POPC lipids in the lower leaflet of the active A_1_AR system were more fluid than in the inactive A_1_AR system. The -S_CD_ order parameters of sn-2 acyl chains of POPC molecules in the upper leaflet were similar in the inactive and active A_1_AR systems. However, the -S_CD_ order parameters for the lower leaflet in the active A_1_AR system were smaller than those in the inactive A_1_AR system. This finding correlated with the outward movement of TM6 in the active A_1_AR, which caused higher inclination of the C-H bonds to be aligned along the bilayer normal. The smaller -S_CD_ order parameters suggested higher membrane fluidity in the lower leaflet of the active A_1_AR system.

In the GaMD simulations, the inactive A_1_AR attracted more lipids in the upper leaflet than the active A_1_AR. The membrane facing positively-charged lysine residue (K265^ECL3^) interacted with the negatively-charged phosphate head group of POPC. In contrast, this lysine pointed towards ECL2 in the active A_1_AR. This was consistent with our previous study^(66)^, in which the positive allosteric modulator (PAM) enhanced the agonist binding at the orthosteric site by forming a salt-bridge between E172^ECL2^-K265^ECL3^. Moreover, the active A_1_AR attracted more lipids in the lower leaflet compared with the inactive A_1_AR. When exposed to the membrane, the positively-charged residues in intracellular region of TM6 of the A_1_AR interacted with negatively-charged head groups. This was further verified by the correlation matrix. The only positive correlation between the transmembrane helices and the lipids was observed between the intracellular region of TM6 and lipids in the lower leaflet of active A_1_AR. Four lysine residues (K263^6.25^, K267^6.29^, K270^6.32^ and K273^6.35^) present in the intracellular end in the β_2_AR of TM6 were also known to interact with negatively-charged headgroups of 1,2-dioleoyl-sn-glycero-3-phosphoethanolamine (DOPE) lipid molecules in and thus stabilize the GPCR active state(27).

In summary, all-atom GaMD simulations have revealed strongly coupled dynamics between a GPCR and the membrane lipids that depend on the receptor activation state. The GaMD method has greatly enhanced sampling of the lipid-protein interactions, which would take significantly longer simulation time using cMD. Nonetheless, the activation or deactivation conformational transitions of the GPCR were not observed in the presented GaMD simulations. Longer GaMD simulations (e.g., microseconds) are expected to capture such conformational transitions (67) and the related effects of lipid-receptor interactions will be investigated in the future. Furthermore, the effects different lipid types (e.g., cholesterol, PIP_2_, etc.) on GPCR-membrane interactions are subject to future studies. It is important to study specific lipid interactions with GPCRs during the receptor activation. Developments of enhanced sampling methodologies and computing power would aid to further address these challenges.

## Acknowledgements

We dedicate this manuscript to Prof. Benoit Roux’s 60^th^ birthday for his contributions in particularly membrane protein simulations and free energy calculations. Computing time was provided on the Comet and Stanford EXtream supercomputers through the Extreme Science and Engineering Discovery Environment award TG-MCB180049 and the Edison and Cori supercomputers through the National Energy Research Scientific Computing Center project M2874. This work was supported in part by the American Heart Association (Award 17SDG33370094), the National Institutes of Health (R01GM132572) and the startup funding in the College of Liberal Arts and Sciences at the University of Kansas.

## Competing Interests Statement

There is no competing interest.

## Graphic Abstract

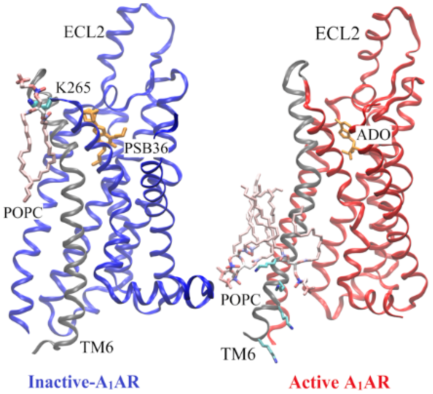
Using adenosine A_1_ receptor (A_1_AR) as a model G-protein-coupled receptor (GPCR), Gaussian accelerated molecular dynamics (GaMD) was applied to explore GPCR-membrane interactions. Membrane lipids played a key role in stabilizing different conformational states of the A_1_AR. Activation of the A_1_AR led to differential dynamics in the upper and lower leaflets of the lipid bilayer. The GaMD simulations revealed strongly coupled dynamics of the GPCR and lipids that depend on the receptor activation state.

