## Supplementary Information for "G-Protein-Coupled Receptor-Membrane Interactions Depend on the Receptor Activation state"

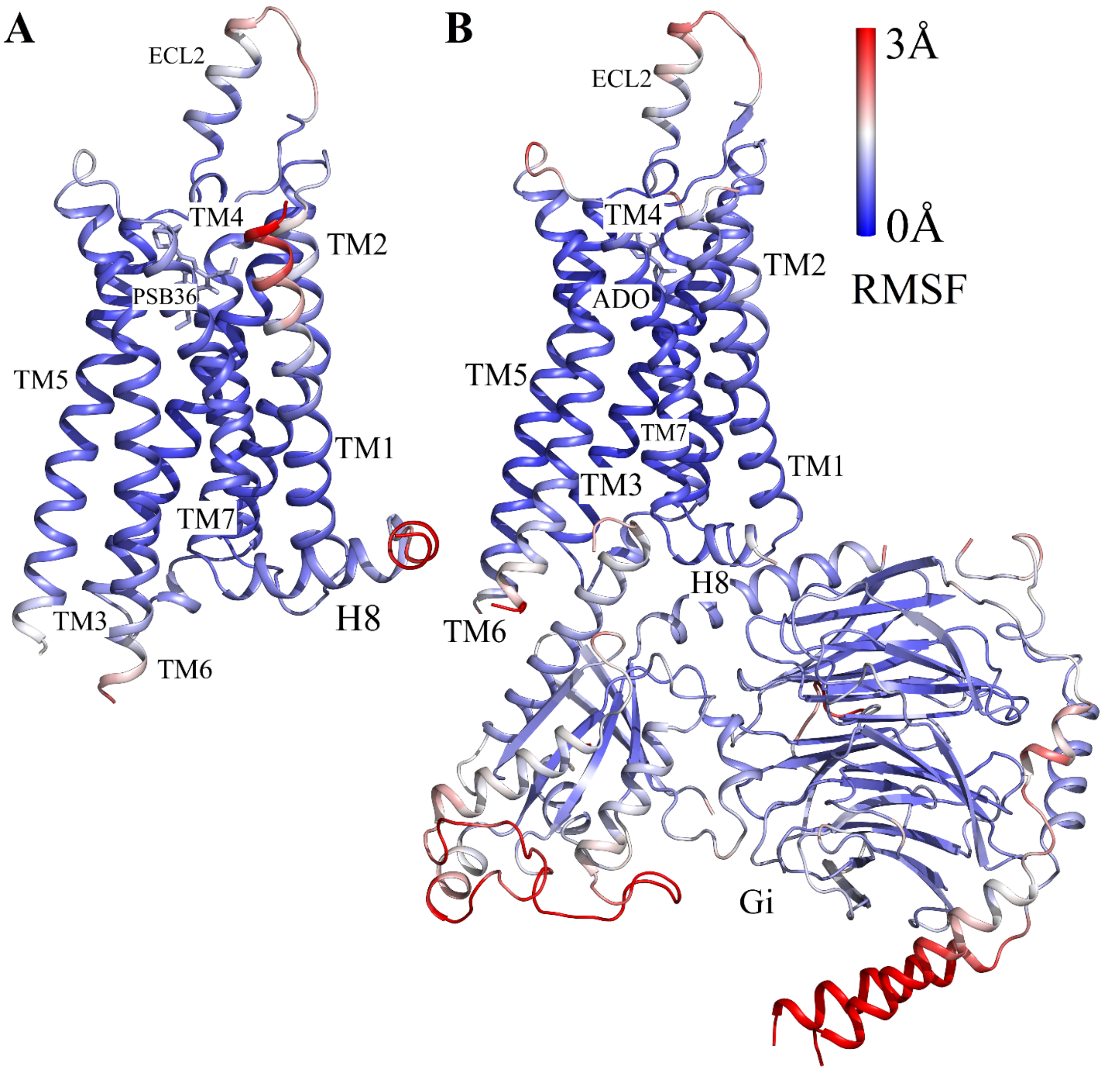


**Figure S1:** Comparison of structural flexibility of the inactive and active $A_1AR$ systems obtained from dual-boost GaMD simulations: (A) Root-mean-square fluctuations (RMSFs) of the inactive PSB36-$A_1AR$ complex. (B) RMSFs of the active ADO-$A_1AR$-Gi complex. A color scale of 0 Å (blue) to 3 Å (red) is used.

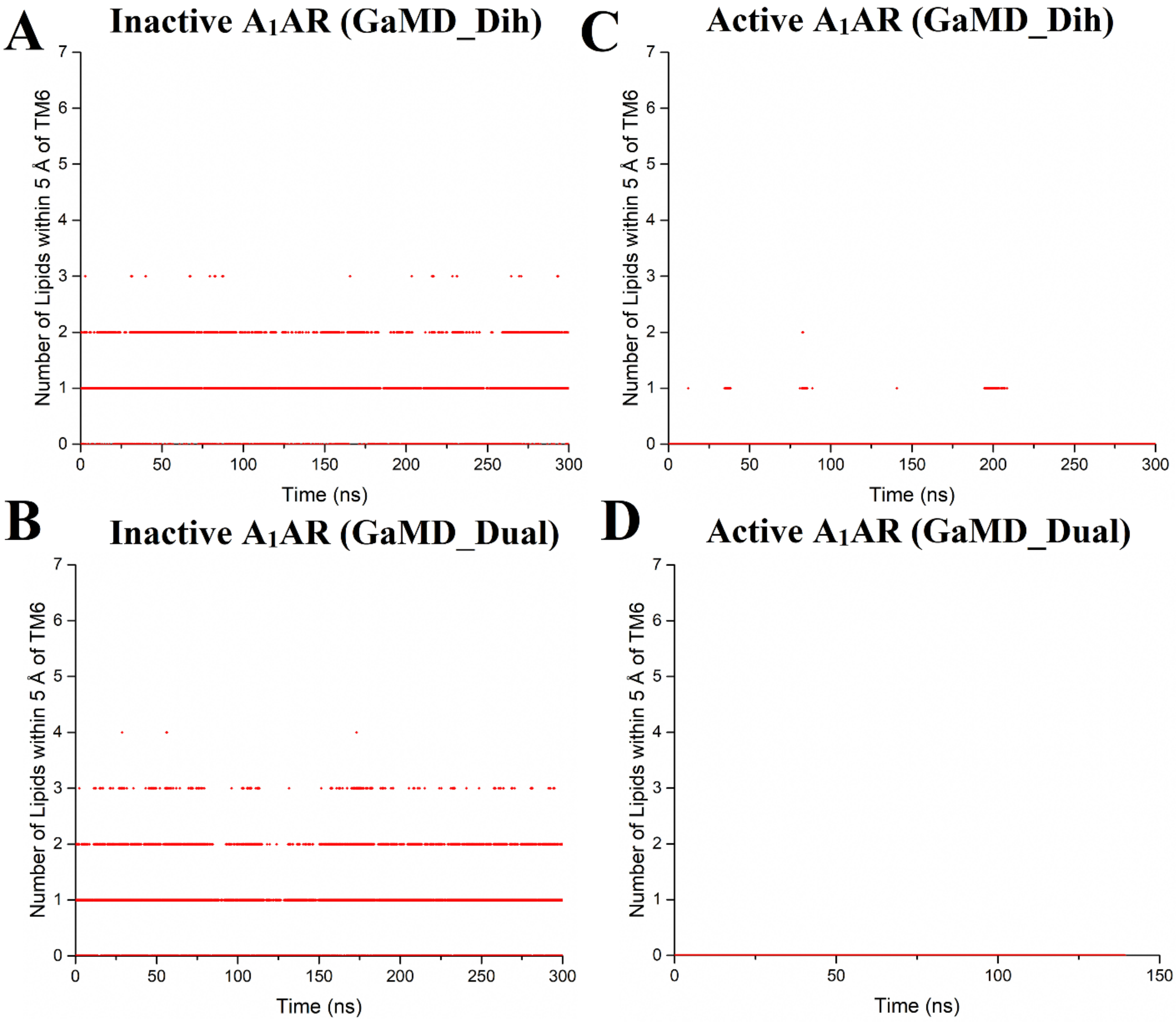


**Figure S2:** Time courses of the number of POPC molecules within 5 Å of TM6 in the upper leaflet of different simulation systems. (A) Inactive $A_1AR$ using dihedral-boost GaMD, (B) Active $A_1AR$ using dihedral-boost GaMD, (C) Inactive $A_1AR$ using dual-boost GaMD and (D) Active $A_1AR$ with using dual-boost GaMD.

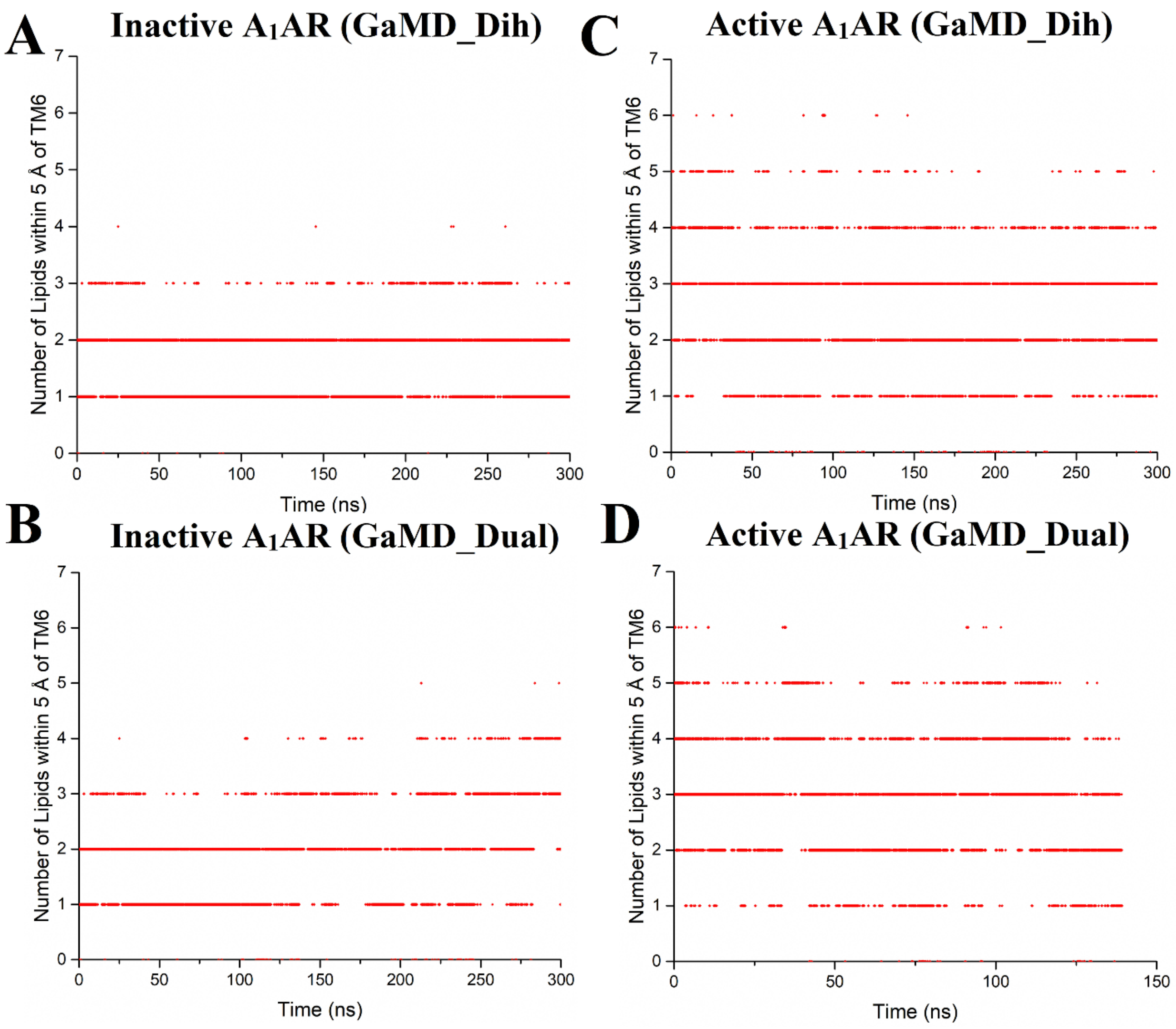


**Figure S3:** Time courses for number of POPC molecules within 5 Å of TM6 in the lower leaflet of different simulation systems. (A) Inactive $A_1AR$ using dihedral-boost GaMD, (B) Active $A_1AR$ using dihedral-boost GaMD, (C) Inactive $A_1AR$ using dual-boost GaMD and (D) Active $A_1AR$ with using dual-boost GaMD.

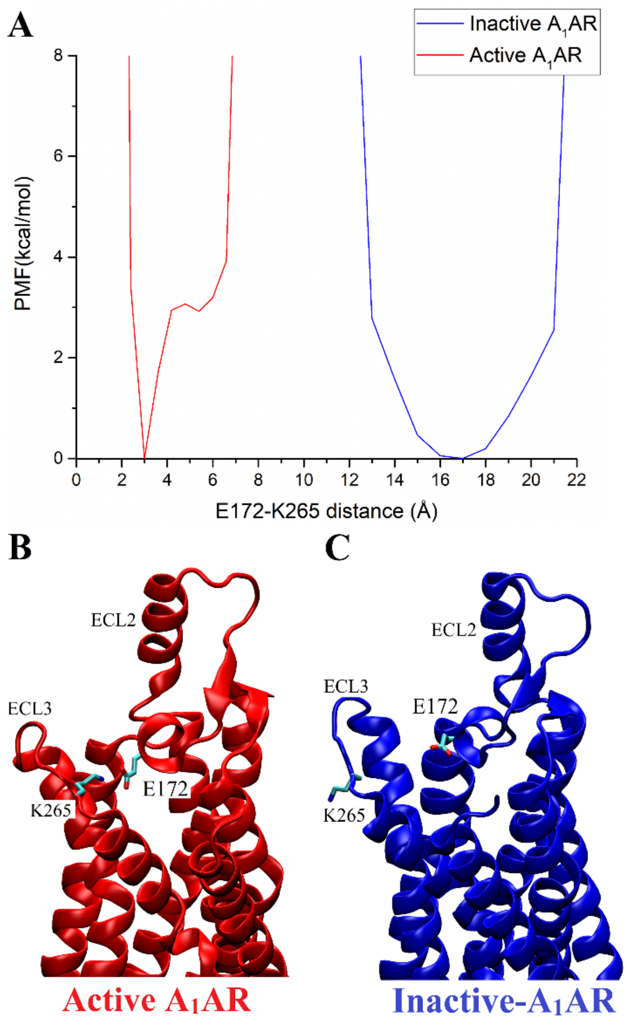


**Figure S4:** Free energy profile of the inactive and active $A_1AR$ systems regarding the $E172^{ECL2}$-$K265^{ECL3}$ distance (A). Minimum energy states of the inactive (C) and active (B) $A_1AR$ systems showing the residues $E172^{ECL2}$ and $K265^{ECL3}$.

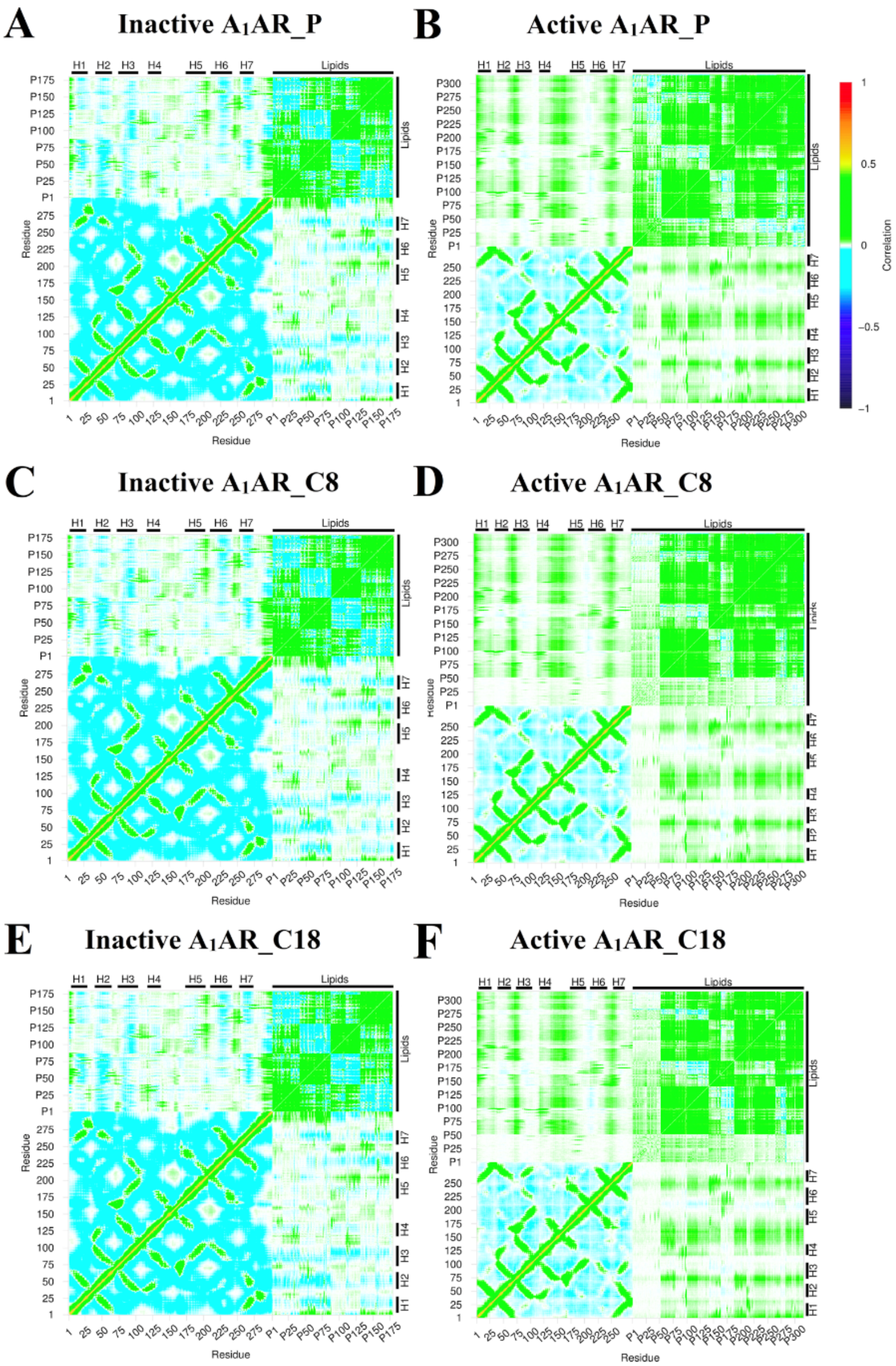


**Figure S5:** Dynamic correlation matrices calculated for lipids with residues in the $A_1AR$ in different simulation systems using dihedral-boost GaMD: (A) Inactive $A_1AR$ using the phosphorous atom in the lipids, (B) Active $A_1AR$ using the phosphorous atom in the lipids, (C)

Inactive $A_1AR$ using the $C_8$ atom in the lipids, (D) Active $A_1AR$ using $C_8$ atom in the lipids, (E) Inactive $A_1AR$ using the $C_{18}$ atom in the lipids, (F) Active $A_1AR$ using the $C_{18}$ atom in the lipids. The $C\alpha$ atoms of the receptor and phosphorous atoms, $C_8$ atoms and $C_{18}$ atoms of POPC lipid were used for calculating the correlation matrices here. These atoms represent different regions of the POPC lipid molecule. Very similar results were obtained using different atoms in POPC lipids to calculate the dynamic correlations with the receptor, similarly for the dual-boost GaMD simulations. The receptor ICL1, ICL2 and ICL3 represent intracellular loops between TM helices 1-2, 3-4, and 5-6 respectively. Similarly, the receptor ECL1, ECL2 and ECL3 represent extracellular loops between TM helices 2-3, 4-5, and 6-7 respectively.
